# Fireworms (*Hermodice carunculata*) are a reservoir and potential vector for coral-infecting apicomplexans

**DOI:** 10.1101/2025.03.06.641876

**Authors:** Anthony M. Bonacolta, Bradley A. Weiler, Candace J. Grimes, Morelia Trznadel, Mark J. A. Vermeij, Patrick J. Keeling, Javier del Campo

## Abstract

Corals (Cnidaria; Anthozoa) play critical roles as habitat-forming species with a wide range, from warm shallow-water tropical coral reefs to cold-water ecosystems [1–3]. They also represent a complex ecosystem as intricate holobionts made up of microbes from all domains of the Tree of Life, that can play significant roles in host health and fitness [4]. The corallicolids are a clade of apicomplexans that infect a wide variety of anthozoans across the world, and can influence the thermal tolerance of habitat-forming corals [1, 5]. Despite their potentially important impacts on reef ecosystems, much of the basic biology and ecology of corallicolids remains unclear. Apicomplexans often have a closed life cycle, with minimal environmental exposure, and sometimes multiple hosts. Corallicolids have only been documented in anthozoan hosts, with no known secondary/reservoir hosts or vectors [6]. Here, we show that abundant corallicolid sequences are recovered from bearded fireworms (*Hermodice carunculata*) in tropical reef habitats off Curaçao, and that they are distinct from corallicolids infecting the corals on which the fireworms were feeding at the time of their collection. The data are consistent with an active infection of fireworms, as opposed to corallicolids being a byproduct of feeding on infected corals, and we suggest that *H. carunculata* is potentially a vector moving corallicolids among coral hosts through its faeces. These findings not only expand our understanding of the ecological interactions within coral reef ecosystems but also highlight the potential role of host-associated parasites in shaping the resilience of reef habitats.

## Main

Multiple *H. carunculata* individuals were observed feeding on the diseased tissue of the boulder star coral, *Orbicella annularis*, at Playa Lagun in western Curaçao (Supplementary Data 1). Triplicate samples (3 samples per subject) were taken of whole worms, a disease transect of *O. annularis* (apparently healthy tissue, diseased transition line tissue, and subsequent necrotic tissue), a healthy *O. annularis* individual from the same reef, a fire worm taken from a healthy *Millepora* sp. colony on a different reef near Piscadera in central Curaçao, and individual water samples from both sampling locations (taken at sampling depth, 1 L water filtered through a 0.2 µm filter). Samples were stored in Zymo DNA/RNA shield before being extracted with the Zymobiomics DNA/RNA Miniprep kit (Catalogue R2002). DNA extracts were PCR-amplified for the V4 regions of 18S rRNA and 16S rRNA genes using established protocols to investigate the eukaryotic nuclear and plastid gene abundances, respectively [7, 8]. Corallicolids, specifically *Anthozoaphila* spp., were found to be the primary microeukaryote recovered from *H. carunculata*, often making up over 50% of the reads (Fig. 1A). Conversely, corals were primarily dominated by Symbiodiniaceae, which gradually decreased in abundance across the disease transect (towards dead tissue). Corallicolids, while still making up a portion of the microeukaryotic community, were not detected in high abundance across the sampled corals, nor in the water in which they were found (Fig 1A).

**Figure 1.**
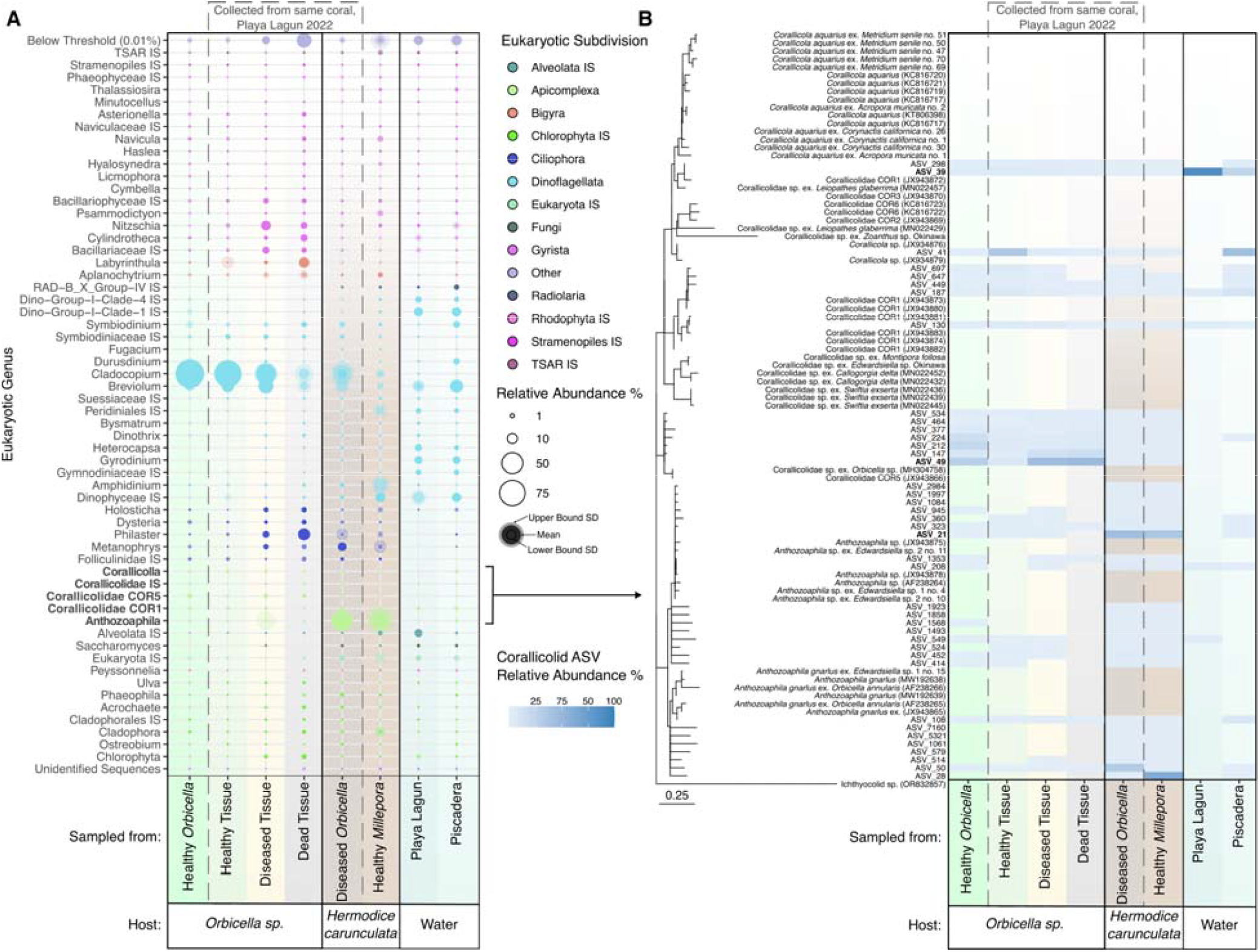
18S rRNA gene metabarcoding data. **(A)** Relative abundance bubble plot of 18S rRNA genes across *Orbicella sp*., *Hermodice carunculata*, and water samples. Bubble shading corresponds to the standard deviation of the samples within each category. The dashed gray box indicates the coral and worm samples taken from the same coral colony in Playa Lagun, Curaçao, in October 2022. **(B)** ASVs classified as corallicolids were EPA-placed onto a backbone 18S-28S rrn operon tree of Corallicolida. Ichthyocolids were used as the outgroup. The relative abundance of each ASV is shown in the heat-map across the same sample categories from panel A.

The primary corallicolid amplicon sequence variants (ASVs) from all samples were placed using RAxML’s Evolutionary Placement Algorithm (EPA) [9] onto an *rrn* operon (18S + 28S rRNA genes) backbone phylogenetic tree to investigate the relatedness of recovered sequences. The predominant corallicolid ASVs from *H. carunculata* were found to branch with *A. gnarlus*, distinctly distant from sequences acquired from the coral on which they were feeding: the coral-derived corallicolids branched with a clade previously recovered from another *Orbicella* sp. (Fig. 1B). The recovery of distinct *A. gnarlus-*related ASVs from multiple *H. carunculata* individuals from multiple locations is notable, as is the absence of corallicolid reads in *H. carunculata* that match those of the disease lesion of the *O. annularis* colony where the worms were feeding. Although corallicolids comprised a tiny portion of the microeukaryotic community in water samples, the predominant corallicolid ASV was again distinct from both the coral and fireworm-associated clades, branching with Corallicolidae COR1 (Fig. 1B).

To better compare our results with other datasets, the plastid 16S rRNA gene was also acquired, as it is a much more common molecular marker in environmental surveys. These data confirmed the high abundance of corallicolids in all *H. carunculata* samples based on plastid 16S rRNA abundance (Fig. 2A), and once again, we also recovered different dominant plastid ASVs for coral, fireworms, and water, respectively, although all ASVs branch together on an EPA-placement tree of previously recovered corallicolid plastid 16S rRNA genes (Fig. 2B).

**Figure 2.**
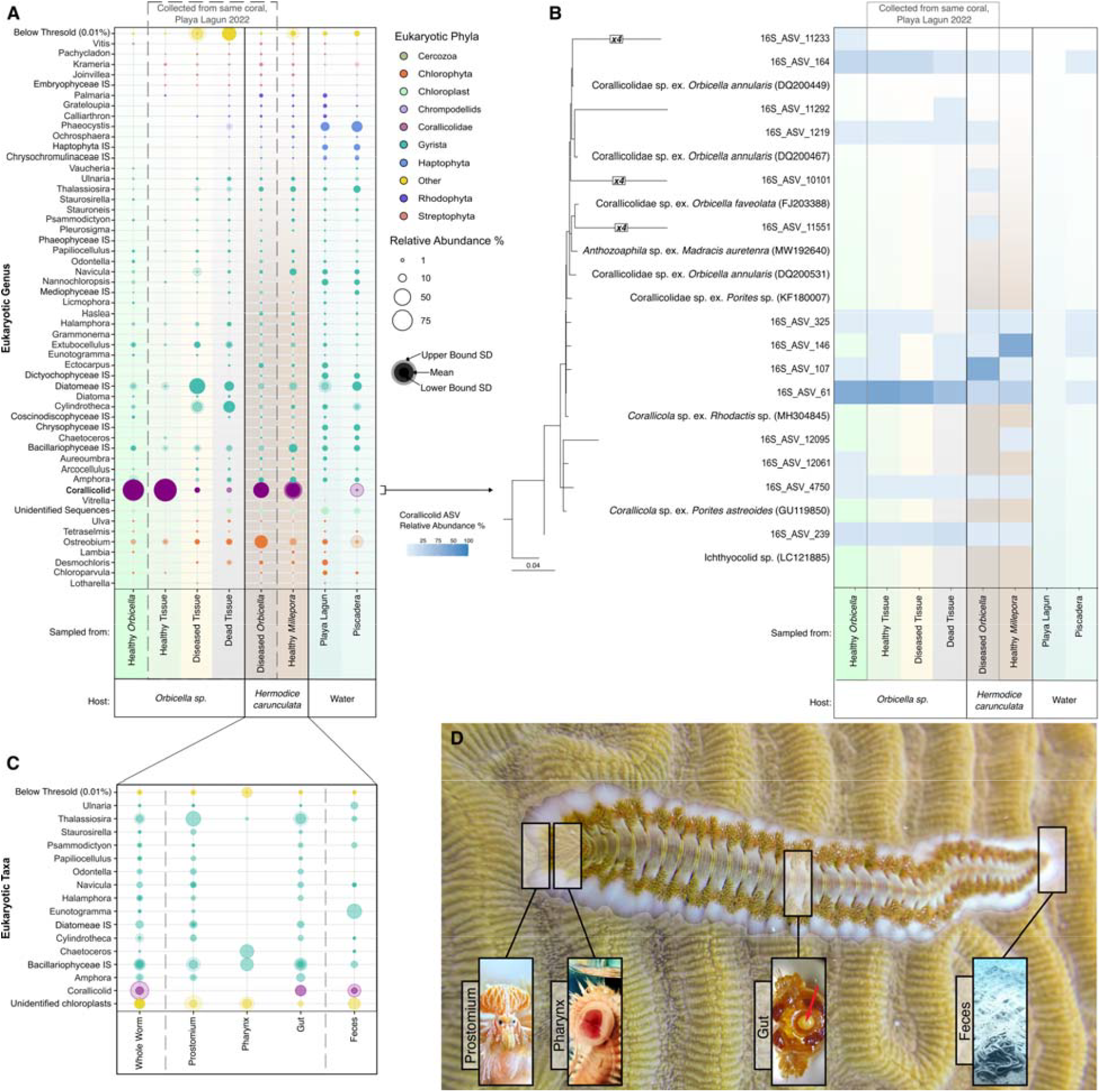
Plastid 16S rRNA gene metabarcoding data. **(A)** Relative abundance bubble plot of plastid 16S rRNA genes across *Orbicella sp*., *Hermodice carunculata*, and water samples. Bubble shading corresponds to the standard deviation of the samples within each category. The dashed gray box indicates the coral and worm samples taken from the same coral colony in Playa Lagun, Curaçao, in October 2022. **(B)** Plastid 16S rRNA ASVs confirmed as corallicolids were EPA-placed onto a backbone 16S rRNA gene tree of Apicomplexa (only the corallicolid portion is shown). The relative abundance of each plastid ASV is shown in the heat-map across the same sample categories from panel A. **(C)** Relative abundance bubble plot of plastid 16S rRNA genes across fire worm sampling sites, indicating the presence of corallicolid ASVs in the whole worm, gut, and feces samples. **(D)** Diagram of *Hermodice carunculata* sampling strategy. Close-up images courtesy of Gabriel Jensen and Candace Grimes.

To examine the tissue specificity in *H. carunculata*, we analyzed 16S rRNA from whole worms and from multiple tissues in dissected *H. carunculata* from Curaçao in 2018. Interestingly, corallicolid plastid 16S rRNA genes were also abundant in the whole worms, as well as the gut and feces, but absent in other tissues (Fig 2C).

Taken together, the data presented here provides the first evidence for a non-anthozoan host of corallicolids. Indeed, the abundance of corallicolids within *H. carunculata* is high compared to the coral specimen. These data also lead to questions of whether *H. carunculata* is infected with a distinct subtype of corallicolid or whether the lifecycle of coral-infecting corallicolids may include other hosts, such as *H. carunculata*. Corallicolids are related to coccidians [10], and many coccidians alternate between sexual reproduction within a primary host and asexual reproduction in a secondary host. The closest sister group to corallicolids is the ichthyocolids, which are blood parasites of marine fishes that are obligately transmitted by gnathiid vectors [11, 12]. It is altogether not unlikely, therefore, that corallicolids also have a life cycle that alternates between hosts. Various apicomplexans also express distinct 18S rRNA genes depending on their life stage (i.e., *Cryptosporidium* [13], *Babesia, Theileria* [14], *Eimeria* [15], and *Plasmodium* [16]). The variation between life-stage 18S rRNAs can be substantial: the 18S rRNAs of the two currently defined corallicolid genera, *Corallicola* and *Anthozoaphila*, share 97.3% identity [6], whereas different life stages of *Plasmodium berghei* can share a lower degree of similarity at 92.0-96.2% [17]. This variation is due to a birth-and-death model of evolution specific to the nuclear rRNA of apicomplexans, likely caused by life stage-specific requirements in rRNA expression patterns [17]. Corallicolid 18S rRNA sequences from coral samples exhibited nearly twice the Shannon-Weiner Alpha diversity compared to fireworm samples (3.35 ± 0.45 vs. 1.78 ± 0.65, p_wilcoxon_ = 1e-07), which may suggest corals are the primary host since sexual and non-sexual life stages are expected to be present concurrently. By contrast, the apicomplexan plastid 16S rRNA gene does not show the same differential expression pattern, and its evolution does not follow the birth-and-death model of its nuclear counterpart. Consistent with this, we find the differences between corallicolid 16S rRNA alpha diversity across coral and fireworm samples are insignificantly different (0.92 ± 0.38 vs. 1.00 ± 0.09, p_wilcoxon_ = 0.82), and indeed we did not observe significant divergence between apicomplexan genera at the 16S rRNA level. While this may be due to life-stage specific expression, nothing is known about the biology of corallicolids, and further investigation would be required to confirm this hypothesis. An alternative hypothesis is that the substitution rate of 18S rRNA is much higher than the 16S rRNA, and the plastid gene accordingly fails to resolve closely-related lineages. It is also possible that the *Anthozoaphila* lineage has an especially wide host range.

The current data suggests corallicolid transmission may be a complex mix of several factors, potentially different in different parasite lineages. For example, brooding coral species have been proposed to transmit corallicolids vertically [18]. Still, some other mode of transmission is required to explain infection in broadcast-spawning coral species, such as *Orbicella*. Fireworms as a vector might fill this gap, as they are common corallivores in tropical reef ecosystems [19] and a fecal-oral transmission mode is consistent with the data. How dynamic this picture might be is exemplified when one also considers that fireworms are invasive in some regions, like the Mediterranean Sea [20]. Also, fireworms have the potential to expand to other ocean regions, such as the Indo-Pacific through the Suez Canal, as a result of the climate crisis and the tropicalization of the Mediterranean Sea [21], so their role as vectors could impact the health of reefs in these regions [22]. And if fireworms are indeed vectors, they cannot be the only species in this role since corallicolids are found in Anthozoans from the deep sea and northern latitudes [2, 3], both outside the range of *H. carunculata* [23], so other, un-explored vectors must operate in these ecosystems. Future research should focus on isolating and sequencing the corallicolids from *H. carunculata* to clarify their association with this host and to expand the search to other potential non-anthozoan hosts to clarify their ecological link to infected anthozoans. Understanding corallicolid ecology in the anthozoan holobiont is critical to uncovering these parasites’ influence in the unprecedented environmental changes these ecosystems are experiencing.

## Data availability

The raw reads for the project have been deposited on NCBI SRA (BioProject: PRJNA1222990). Code used for analysis and ASV count tables, taxonomy tables, and nucleotide sequences can be found on GitHub at: https://github.com/delCampoLab/Hermodice_corallicolids/.

## Acknowledgments

We thank Gabriel Jensen (Instagram: @shallowseasgallery) for the photographs. We also acknowledge the University of Miami start-up funding, project PID2020-118836GA-I00 financed by MICIU/AEI/10.13039/501100011033, and project 2021 SGR 00420 financed by the Departament de Recerca i Universitats de la Generalitat de Catalunya for JDC, University of Miami IDSC Resources for Early Career Researchers for BAW, and Texas Sea Grant’s Grants-in-Aid of Graduate Research Program (FY2019-2021) and the Galveston Graduate Student Association for CG. All sample exports were conducted under CITES Permits #20US835702/9 & 2012/48584. Further, we acknowledge Dr. Anja Schulze at Texas A&M University Galveston and Dr. Jose V. Lopez at Nova Southeastern University for assistance with CJG dissertation research and collection of the 2018 samples.

## Notes

https://github.com/delCampoLab/Hermodice_corallicolids/tree/main

### Competing Interest Statement

The authors have declared no competing interest.

