## Supplementary figures and images for "Fireworms (*Hermodice carunculata*) are a reservoir and potential vector for coral-infecting apicomplexans"

### Supplementary Data 1

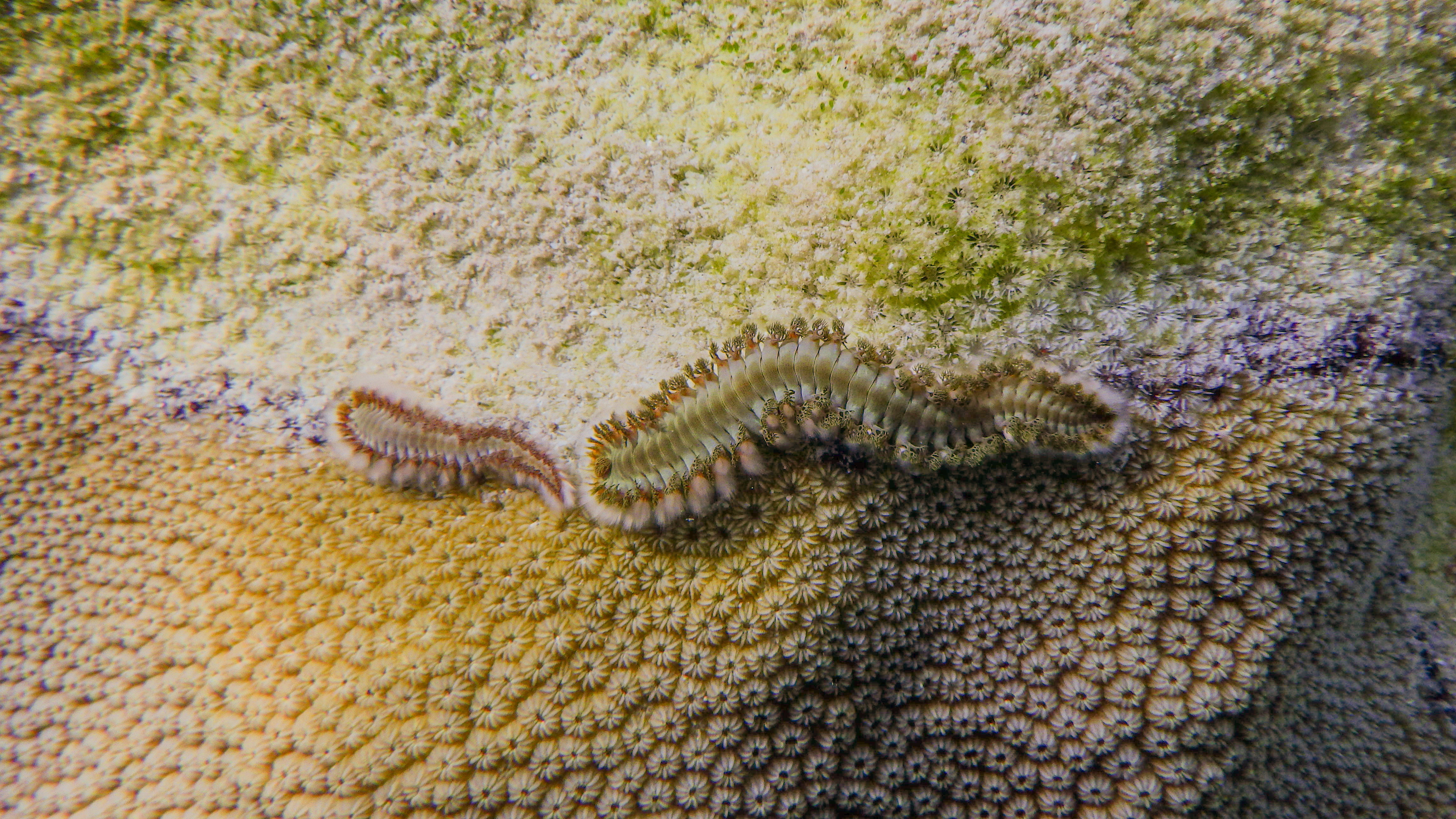
